# The Senescence-Associated Secretory Phenotype constitutes HIF-1α activation but is independent of micronuclei-induced cGas/Sting activation

**DOI:** 10.1101/2024.09.25.615081

**Authors:** Celestine Z. Ho, Lin Deng, Remigio Picone, Fieda Abderazzaq, Nicole Flanagan, Dominic Zhuohong Chua, Boon Chuan Low, Selwin K. Wu

**Author notes:** Correspondence, Present address: Mechanobiology Institute, National University of Singapore, Singapore. Equal contributions (Celestine Z. Ho, Lin Deng).

## Abstract

The Senescence-Associated Secretory Phenotype (SASP), characterized by the upregulation of inflammatory cytokines, is triggered during senescence by anti-proliferation stresses, including replicative exhaustion, γ-irradiation, Ras oncogene induction, and centrosome amplification. The elucidation of common signalling pathway(s) activated in SASP, induced by different anti-proliferation stresses, remains an important question. Indeed, micronuclei activation of the cGAS/Sting pathway, which has been thought to drive SASP(Kwon, Leibowitz, and Lee, 2020), remains controversial(Flynn, Koch, and Mitchison, 2021; Sato and Hayashi, 2024; Takaki *et al*., 2024). In this report, analyses of various cell lines induced to undergo senescence by diverse stressors revealed that HIF-1α is specifically induced in senescence but not in quiescence. Consistent with our previous findings(Wu *et al*., 2023a), we have further demonstrated how centrosome amplification induces a non-canonical SASP dominated by HIF-1α activation rather than the classical NFκB signaling. Lastly, we revealed that during SASP, centrosome amplification-generated micronuclei do not activate the cGAS/Sting-mediated interferon response. Together, our findings demonstrate that HIF-1α-activation in SASP is a defining feature of the SASP induced by diverse stressors, acting independently of micronuclei generation and cGAS/Sting activation.

## INTRODUCTION

Cellular senescence is characterized by the upregulation of pro-inflammatory cytokines, a phenomenon termed the Senescence-Associated-Secretory Phenotype (SASP)(Coppe *et al*., 2010). Senescence is triggered by many stresses(Ben-Porath and Weinberg, 2004), including γ-irradiation, oncogene induction, replicative exhaustion(Purcell, Kruger, and Tainsky, 2014), and centrosome amplification(Wu *et al*., 2023a). The elucidation of common signalling pathway(s) activated in SASP, induced by different anti-proliferation stresses, remains an important question. Here, we first provide a background on the consequences of centrosome amplification. Then, we focus on centrosome amplification, generating micronuclei and their controversies with Cyclic guanosine Monophosphate (GMP)-AMP synthase (cGAS)/Stimulator of interferon genes (STING) activation. Lastly, we delve into the background of another pathway that participates in SASP, the hypoxia-inducible factor-1α (HIF-1α) pathway. We test the hypothesis that SASP constitutes HIF-1α activation but is independent of micronuclei-induced cGas/Sting activation.

Centrosomes are microtubule-organizing centers that support cellular processes such as cell division and adhesion(Akhmanova, Stehbens, and Yap, 2009). In most normal cells, centrosomes are duplicated once during each cell division, such that G1 phase cells typically have one centrosome and mitotic cells have two. In contrast, senescent and cancer cells commonly exhibit increased numbers of centrosomes, termed centrosome amplification. Centrosome amplification can induce proliferation arrest by the PIDDosome multiprotein complex(Fava *et al*., 2017) and Hippo pathway-mediated mechanisms(Wu *et al*., 2023a). Centrosome amplification also stimulates reactive oxygen species production(Wu *et al*., 2023a) to upregulate the expression of SASP factors. These secreted SASP factors induce cell-cell contact disruption(Mangold *et al*., 2011), tumor growth(Angelini *et al*., 2013), and invasion(Wu *et al*., 2023a).

Besides the induction of SASP through ROS in centrosome amplification(Wu *et al*., 2023a), centrosome amplification also causes chromosome missegregation and micronuclei formation, leading to aneuploidy(Ganem, Godinho, and Pellman, 2009). Micronuclei, a major source of aneuploidy, have been reported to activate the cGAS/STING pathway with γ-irradiation or MPS1 inhibition (for a longer duration and at a higher dose than our MPS1i treatment)(Harding *et al*., 2017; Mackenzie *et al*., 2017; Mohr *et al*., 2021). In these cases, cGAS/STING activation was shown to be essential for SASP(Gluck *et al*., 2017). How is cGAS/STING activated? cGAS is a cytosolic DNA sensor that triggers the innate immune interferon response when it binds to dsDNA in the cytosol(Li *et al*., 2013). Briefly, activated cGAS, when bound to dsDNA, converts ATP and GTP into 2′3′-cyclic GMP-AMP (cGAMP). cGAMP then binds to and activates STING, leading to TANK-binding kinase 1-dependent phosphorylation, nuclear translocation of interferon regulatory factor 3, and transcriptional activation of interferon-stimulated genes(Ablasser and Hur, 2020).

Senescence-inducing triggers such as γ-irradiation or MPS1 inhibition (at a longer duration with a higher dose of inhibition than the MPS1i treatment used here) generate micronuclei with defective nuclear envelopes, exposing micronuclei DNA to the cytoplasm, leading to cGAS/Sting activation(Harding *et al*., 2017; Mackenzie *et al*., 2017; Mohr *et al*., 2021). Despite these studies, the role of micronuclei in activating cGAS/STING remains controversial. Thus, we next highlight studies indicating that micronuclei do not activate cGAS/STING(Flynn, Koch, and Mitchison, 2021; Sato and Hayashi, 2024; Takaki *et al*., 2024).

Recent reports with a stronger focus on the analysis of individual cells(Flynn, Koch, and Mitchison, 2021; Sato and Hayashi, 2024; Takaki *et al*., 2024) indicate that micronuclei from chromosome missegregation fail to activate cGAS/Sting-mediated innate immune response(Flynn, Koch, and Mitchison, 2021; Sato and Hayashi, 2024; Takaki *et al*., 2024). cGAS localization to nuclear envelope-ruptured micronuclei during interphase was reported to be rare(Flynn, Koch, and Mitchison, 2021; Sato and Hayashi, 2024; Takaki *et al*., 2024), with cGAS more often binding to micronuclei during mitosis and remaining on cytosolic chromatin. Notably, cGAS accumulation during mitosis did not activate STING or trigger the interferon response in the subsequent interphase(Sato and Hayashi, 2024; Takaki *et al*., 2024). Amongst the cellular senescence triggers, γ-irradiation activated STING independently of micronuclei formation and cGAS localization to micronuclei, suggesting that micronuclei do not trigger cGAS/STING pathway activation(Takaki *et al*., 2024). Indeed, STING-mediated interferon response after γ-irradiation is linked to cytosolic mitochondrial DNA release instead(Sato and Hayashi, 2024). The presence of histones in micronuclei was found to be a potential inhibitory factor, preventing cGAS activation(Takaki *et al*., 2024).

Besides the controversies surrounding cGAS/Sting activation by micronuclei, we next focus on another critical pathway participating in SASP: hypoxia signaling and HIF-1α activation(Wu *et al*., 2023a). We have discovered that HIF-1α is activated in centrosome amplification, oncogene, and γ-irradiation-induced senescence(Wu *et al*., 2023a). Hypoxia-inducible factors are key transcription factors that regulate oxygen sensing and mediate the hypoxic response(Kietzmann, Mennerich, and Dimova, 2016). Activation of hypoxia-inducible factors triggers the transcription of genes crucially involved in cancer(Miroshnikova *et al*., 2016) and immunity(Palazon *et al*., 2014), including cytokines such as IL8, IL6, ANGPTL4, VEGF, and PDGF, all commonly observed in SASP (Yoshida *et al*., 2006; Maxwell *et al*., 2007; Liu *et al*., 2009; Coppe *et al*., 2010).

Cellular senescence triggers, such as constitutively active Ras overexpression, which promotes cell invasion(Wu *et al*., 2014b; Wu *et al*., 2015) (Wu and Yap, 2013), are known to activate HIF-1α(Mazure *et al*., 1996; Mazure *et al*., 1997; Wu *et al*., 2023a). γ-irradiation, another senescence trigger, is also known to trigger HIF-1α activation(Moeller *et al*., 2004; Wu *et al*., 2023a). However, reports indicate that hypoxia and HIF-1α activation can prevent or delay cells from entering senescence (Welford and Giaccia, 2011). Nevertheless, the HIF-1α activity of cells once they have entered replicative exhaustion-induced senescence has not been studied.

In this report, we reveal that HIF-1α target genes are specifically induced in senescence but not in the reversible cell cycle arrest-quiescence. We then focused on centrosome amplification-induced senescence. Consistent with our previous finding(Wu *et al*., 2023a), we further demonstrate that centrosome amplification induces a non-canonical SASP dominated by HIF-1α activation. Finally, we show that during SASP, centrosome amplification-generated micronuclei do not activate the cGAS/Sting-mediated interferon response. In all, we conclude that SASP constitutes HIF-1α activation but is independent of the micronuclei-induced cGas activation.

## MATERIALS & METHODS

### Cell culture

To prevent mycoplasma contamination, all source cell cultures were maintained in media containing plasmocin (1:5000 dilution, Invivogen). Culture plates were briefly coated with 40μg/ml of fibronectin (Sigma)(Ho *et al*., 2025), and glass coverslips were incubated in 40 μg/ml fibronectin for at least 2 hours. Human mammary epithelial MCF10A cells were maintained at 37^0^C with 5% CO2 atmosphere and cultured as previously described^14^. In brief, MCF10A cells were cultured in phenol red-free DMEM/F12 (Invitrogen) supplemented with 5% horse serum (Sigma), 20ng/ml epidermal growth factor (EGF; Sigma), 10mg/ml insulin (Invitrogen), 100 mg/ml hydrocortisone (Sigma), 1ng/ml cholera toxin (Sigma) and 100 IU/ml penicillin and 100 μg/ml streptomycin (Invitrogen). WI38 lung fibroblast cells, obtained from the ATCC, were cultured in ATCC-formulated Eagle’s Minimum Essential Medium containing 10% FBS. Telomerase-immortalized RPE-1 cells (ATCC) and all derivative cell lines generated in this study were cultured in phenol red-free DMEM/F12 media containing 10% FBS, 100U/ml penicillin, and 100μg/ml streptomycin. RPE-1 cells were maintained at 37°C in 5% CO2 atmosphere.

To conditionally overexpress PLK4, lentiviral vectors pLenti-CMV-TetR-Blast (17492, Addgene) and pLenti-CMV/TO-Neo-Dest (17292, Addgene) were used. Wild-type PLK4 and PLK4 1–608 cDNAs (control) were cloned into the pLenti-CMV/TO-Neo-Dest vector using the Gateway system. W138 lung fibroblasts were coinfected with a lentivirus containing the TetR and the lentivirus containing the wild-type PLK4 or control PLK4^1-608^ open reading frames.

### Induction of centrosome amplification and senescence

At day 0, control PLK4^1-608^ and PLK4 cells were gently dissociated into single cells, then seeded at low density into media with 2μg/ml of doxycycline such that they were ∼80% confluent on day 5. After ∼48 hours of doxycycline incubation, the culture media with doxycycline was replaced. On day 5, cells are gently dissociated into single cells before reseeding into 6 well plates to ∼80% confluent on day 7. Because Control PLK4^1-608^ cells grow faster than PLK4-induced cells, Control PLK4^1-608^ cells are seeded at ∼2.5 times lower density than PLK4-induced cells. Cells at ∼80% confluency are analyzed on day 7 or 8. Media is left unchanged for ∼3 days before analysis.

### Generation of tetraploids and ‘evolved’ tetraploids with normal centrosome numbers

To generate G1 arrested tetraploids, 15cm dishes were seeded with 6 million exponentially growing RPE1-FUCCI cells, such that they were ∼65% confluent the following day. Day 2: 4 μM Dihydrocytochalasin B (DCB) was added to each 15 cm dish for 16 hr. Day 3: DCB-treated cells were washed 5 times for 5 minutes, incubated in medium containing 2.5 μg/ml Hoechst dye for 1 hr, then trypsinized in 0.05% trypsin. G1 diploids (2C DNA content, C refers to the DNA content measured by the FACs, mCherry+, GFP−) and G1 tetraploids (4C DNA content; mCherry+, GFP−) were isolated by FACS sorting. FACS-sorted cells were seeded on 6 well plates coated with 40μg/ml fibronectin.

Evolved tetraploid RPE-1 cells, kindly provided by Huadi Zhang, with the normal number of centrosomes were derived as described(Ganem, Godinho, and Pellman, 2009). Briefly, tetraploid RPE-1 cells were treated for 16 hr with 0.2 μM cytochalasin D and FACS-sorted by DNA content using Hoechst staining (Molecular Probes). The peak of cells with a DNA content of 8C were isolated and cultured for ∼1 week before a second iteration of the FACS sorting to re-isolate 8C cells. A total of 5 purifying cell sortings for ∼6 weeks were required to generate a pure population of proliferating tetraploid cells with normal centrosome numbers.

### CRISPR Knockouts and Lentiviral Transduction

The previously described(Cho *et al*., 2016) HIF-1α pLenti-CRISPRv2 was a gift from William G. Kaelin, Jr. (Dana-Farber Cancer Institute, Boston, USA). The sgRNA sequence targeting HIF-1α is ‘5’-CACCGTGTGAGTTCGCATCTTGATA-’3’. Clonal selection for single-cell clones after puromycin selection was used for the experiments.

For lentivirus production, 6μg lentiviral constructs, 3 μg psPAX2 (Addgene), and 3 μg pMD2.G (Addgene) were transfected into 293T cells at 80% confluency in a 10-cm dish with 8 ml of media. 20 μl of lipofectamine 3000 (Life Technologies) was used and transfection was performed according to manufacturer’s instructions. Virus was harvested at 48 hours and 72 hours post-infection and filtered with a 0.2μm filter(Smutny *et al*., 2011; Moore *et al*., 2014). Cells were infected for 12 hours, with virus containing the genes of interest in the presence of 10 μg/ml polybrene, washed thoroughly, and allowed to recover for 24 hours before any selection(Smutny *et al*., 2011; Moore *et al*., 2014).

### Antibodies and drugs

Primary antibodies used in this study were: rabbit monoclonal antibodies against cGAS (1:1000 for western blotting; 1:200 for immunofluorescence; Cell Signaling Technologies; catalogue number 15102S, clone number D1D3G), rabbit polyclonal antibodies against HIF-1α overnight at 4 degrees Celsius (1:50-1:100 for immunofluorescence; Genetex; catalogue number GTX127309), rabbit polyclonal antibodies against phospho-STING Ser366 (1:200 for immunofluorescence; Cell Signaling Technology; catalogue number 13647), rabbit monoclonal antibodies against Ki-67 (1:200 for immunofluorescence; Invitrogen; catalogue number MA5-14520, clone number SP6), mouse monoclonal antibodies against Centrin (1:200 for immunofluorescence; MilliporeSigma; catalogue number 04-1624), mouse monoclonal antibodies against HP1γ (1:200 for immunofluorescence; MilliporeSigma; catalogue number MABE656, clone number 14D3.1), rabbit polyclonal antibodies against Laminin overnight at 4 degrees Celsius (1:200 for immunofluorescence; Novus Biologicals; catalogue number NB300-144); mouse monoclonal antibody against GAPDH (1:1000 for western blotting; Santa Cruz Biotechnology; catalogue number sc-32233), Alexa 647-conjugated phalloidin (1:400 for immunofluorescence; Life Technologies; catalogue number A22287).

Doxycycline (Sigma Aldrich catalogue number D3072) was used at 2 mg/ml, and TNFα (Roche Diagnostics catalogue number 11371843001) was used at 20ng/ml for 2 hours. The following inhibitor doses were used: 0.1 and 0.2 μM Reversine for 24 hours (Cayman Chemical) 4μM dihydro cytochalasin B (DCB; Sigma).

### Immunofluorescence and live-cell microscopy

Cells were fixed at 4°C with 4% paraformaldehyde in cytoskeleton stabilization buffer (10mM PIPES at pH 6.8, 100mM KCl, 300mM sucrose, 2mM EGTA, and 2mM MgCl2) on ice for 25 min and subsequently permeabilized with 0.2% Triton-X in PBS for 10 minutes at room temperature(Smutny *et al*., 2011; Han *et al*., 2014; Leerberg *et al*., 2014; Wu *et al*., 2014a; Gomez *et al*., 2015; Acharya *et al*., 2017; Wu *et al*., 2023b). Fixed cells were blocked with 5% BSA in TBS overnight at 4°C. Primary antibodies were incubated overnight at 4°C for HIF-1α and 45 minutes at room temperature for p65/RELA. For all immunostaining, secondary Alexa Fluor antibodies were incubated at room temperature for an hour(Mangold *et al*., 2011; Han *et al*., 2014; Leerberg *et al*., 2014; Gomez *et al*., 2015; Greenlees *et al*., 2015; Acharya *et al*., 2017). For live cell imaging, time-lapse images were collected with a Nikon Ti inverted microscope with 20x wide-field optics equipped with a perfect focus system, a Hamamatsu ORCA ER cooled CCD camera controlled with Nikon NIS-Element software, and an OkoLab 37°C, 5% CO_2_ microscope chamber. Confocal images were also acquired using a Yokogawa CSU-W1 Spinning Disk microscope (Nikon) equipped with a 40× 0.95 NA objective or 60× 1.20 NA water immersion objective or 100× 1.45 NA oil immersion objective.

### Image processing and analysis

ImageJ macros are created to determine the HIF-1α nuclear and cytoplasmic fluorescence intensity. First, we determine the nuclear-to-cytoplasmic intensity ratio for HIF-1α. We have previously determined that the cytoplasmic HIF-1α intensity is largely background intensity unaffected by HIF-1α knockout. Therefore, we subtract a value of 1 from the HIF-1α nuclear-to-cytoplasmic intensity ratio, as we consider any HIF-1α nuclear intensity equal to the cytoplasmic intensity as background. For the final subtracted value, any negative values are considered zero for HIF-1α nuclear levels.

To quantify the nuclear and cytoplasmic mean fluorescence intensity of HIF-1α and p65, a nuclei mask was generated from the Z-projection of the Hoescht nuclear staining. A 1 μm region surrounding the nucleus was designated as the cytoplasmic region. The watershed function was applied to the nuclei mask to ensure the separation of overlapping nuclei.

### β-Galactosidase Staining

The scoring of SA-β-gal positive cells was determined using the Senescent Cell Staining Kit (Sigma) according to the manufacturer’s instructions. The percentage of positive cells was counted manually on a microscope with a bright lamp and a color camera to capture the blue stain. We find that the Senescent Cell Staining Kit from Sigma is more sensitive and thus more reliable for detecting centrosome amplification-induced senescence than the Abcam kit.

### Media Swap Motility Assay

Cells were plated at 0.5-1×10^6^ per well in 6 well plates and allowed to attach for 24h. Then, cells were starved in serum-free medium for two hours and incubated in 0.5% horse serum media for another 12 hours. Conditioned medium from cells with or without centrosome amplification were collected and centrifuged at 5000 rpm for 20 minutes to eliminate detached cells or debris. Conditioned medium was then added to cells, and the effect of the swapped conditioned medium on cellular motility was imaged with bright field microscopy (Nikon Ti).

### Paracrine Factors Induced Co-culture Invasion Assay

Cells were grown in the same medium with horse serum (5%) and EGF (20ng/ml). To assay invasion in 3D cultures, MCF10A cells were trypsinized into single cells and seeded in the 12mm well of a 35mm glass base dish (Iwaki) plated with a 200μl mix of Matrigel (Corning) and 7% collagen-I (Gibco) to facilitate invasion. Media used in 3D cultures was supplemented with reduced horse serum (2%), EGF (20ng/ml), and Matrigel (2%). Control or centrosome amplification-induced senescence PLK4 cells were seeded at the fibronectin-coated polystyrene sides of the glass base dish in media supplemented with horse serum (5%) to form a confluent monolayer, ensuring that both cultures avoid contact. After cell attachment, cells are the sides of the dish were washed with PBS, and 2ml media was added to the co-culture setup. The assay was kept in culture at 37°C for 4 days prior to live imaging and quantification of invasive structures.

### ELISA

For DMOG treatment, cells were seeded with 1μm DMOG and incubated overnight at 37°C. On day 5, cells for all conditions were starved with serum-free media for 2 h and replaced with full media for 24h. Conditioned media was subsequently collected and supplemented with (concentration) protease inhibitor cocktail. Media was centrifuged at 5000 rpm for 10 minutes, and supernatant was collected. ELISA was subsequently performed using Human Cytokine Antibody Array (Abcam) according to the manufacturer’s instructions.

### NextSeq500 RNA Library Preparation and Sequencing

RNA quantity was determined on the Qubit using the Qubit RNA Assay Kit (Life Tech), and RNA quality was determined on the Bioanalyzer using the RNA Pico Kit (Agilent). Using the NEB Next Ultra RNA Library Prep Kit for Illumina (NEB), which selects for poly(A) mRNA, 100ng of total RNA was used for cDNA library construction, according to the manufacturer’s protocol. Following library construction, DNA quantity was determined using the Qubit High Sensitivity DNA Kit (Life Technologies), and library insert size was determined using the Bioanalyzer High Sensitivity Chip Kit (Agilent). Finally, qPCR was performed using the Universal Library Quantification Kit for Illumina (Kapa Biosystems) and run on the 7900HT Fast qPCR machine (ABI) to quantify and normalize libraries. Libraries that passed quality control were diluted to 2nM DNA content using sterile water and then sequenced on the NextSeq500 (Illumina) at a final concentration of 12pM, according to the manufacturer’s instructions.

### RNA Sequencing Analysis

Sequencing reads were aligned to the reference genome using the RNA-specific STAR aligner using default parameters(Dobin *et al*., 2013). The quality of sequencing runs and alignments was assessed by RNA-SeQC(DeLuca *et al*., 2012). The featureCounts tool was used to count sequencing reads mapped to the reference genome at the gene level(Liao, Smyth, and Shi, 2014). The counts were subsequently normalized between samples using DESeq2 to control for experimental variation(Anders and Huber, 2010). Differential expression analysis was performed between the desired experimental contrasts using DESeq2.

### Gene Set Enrichment Analysis

We used GSEA v3.0 to examine the association between gene sets and gene expression. Normalized read counts from DESeq2 analysis were used for GSEA. Genes with a 0 read count in either control or experimental group were excluded. GSEA was performed using gene sets as permutation type, 1,000 permutations, and log2 ratio of classes as a metric for ranking genes. To construct a HIF-1α signature up regulated gene set, we performed RNA sequencing of MCF10A cells after overnight treatment with 1 μM of DMOG, with DMSO-treated cells as control. We included genes that were upregulated by DMOG in DESeq analysis (*p*(adjusted) < 0.05, fold change ≥4). To construct a DNA-damage senescence gene set, we performed RNA sequencing of MCF10A cells 7 days after exposure to 12Gy of γ-irradiation. We obtained published data sets to construct both upregulated and downregulated core senescence gene sets(Hernandez-Segura *et al*., 2017).

### Quantitative RT-PCR

Total RNA was isolated using the RNAeasy Mini Kit (Qiagen), and cDNA was synthesized using SuperScript III (Invitrogen) according to manufacturer instructions. Quantitative RT-qPCR was performed in triplicate using SYBR Green (Life Technologies) on a ViiA 7 Real-Time PCR machine (Thermo Fisher Scientific) with GAPDH as an internal normalization control. Sequences of qPCR primers for cGAS are as follows: Forward primer: 5’-gggagccctgctgtaacacttcttat-3’, Reverse primer: 5’-cctttgcatgcttgggtacaaggt-3’

## RESULTS AND DISCUSSION

### The hallmark hypoxia genes are induced in cellular senescence

We began by analyzing the gene expression signature activated in cellular senescence using existing databases and from our experiments. To confirm that these expression data sets are from cells that are in senescence, we validate the datasets by analyzing their senescence expression pattern. Despite variabilities in the senescent phenotype(Sharpless and Sherr, 2015; Hoare *et al*., 2016; Schmitt, 2016; Lee and Schmitt, 2019), a core transcriptome signature of a list of upregulated and downregulated genes has been identified as a useful marker to consistently identify senescent cells(Hernandez-Segura *et al*., 2017). Thus, we employ this universal senescent gene set to confirm the senescent identity of cells that underwent γ-irradiation, replicative exhaustion, and constitutive Ras oncogene-induced senescence. MCF10A untransformed immortal breast epithelial cells were exposed to 12 Gy of γ-irradiation, whereas the replicative exhaustion(Kang *et al*., 2015) and constitutive active Ras-induced IMR90 normal fibroblasts(Acosta *et al*., 2013) transcriptomes were obtained from published datasets(Kang *et al*., 2015) investigating the senescence-associated secretory phenotype. The IMR90 fibroblasts analyzed are commonly used to study senescence, as IMR90 can undergo replicative exhaustion-induced senescence after ∼58 population doublings. Seven days after irradiation, the transcriptomes of control and γ-irradiated cells were compared. GSEA determines the transcriptional program activated by identifying coordinated changes in a defined set of gene expressions(Subramanian *et al*., 2005). Gene set enrichment analysis revealed a strong association of γ-irradiation (Fig. 1a,b), replicative exhaustion (Fig. 1c,d), and constitutive-active Ras oncogene (Fig. 1e,f) induced genes with both upregulated and downregulated senescence gene sets. Thus, this provides evidence that these transcriptomes are from senescent cells generated by different senescence-inducing stresses.

**Figure 1.**
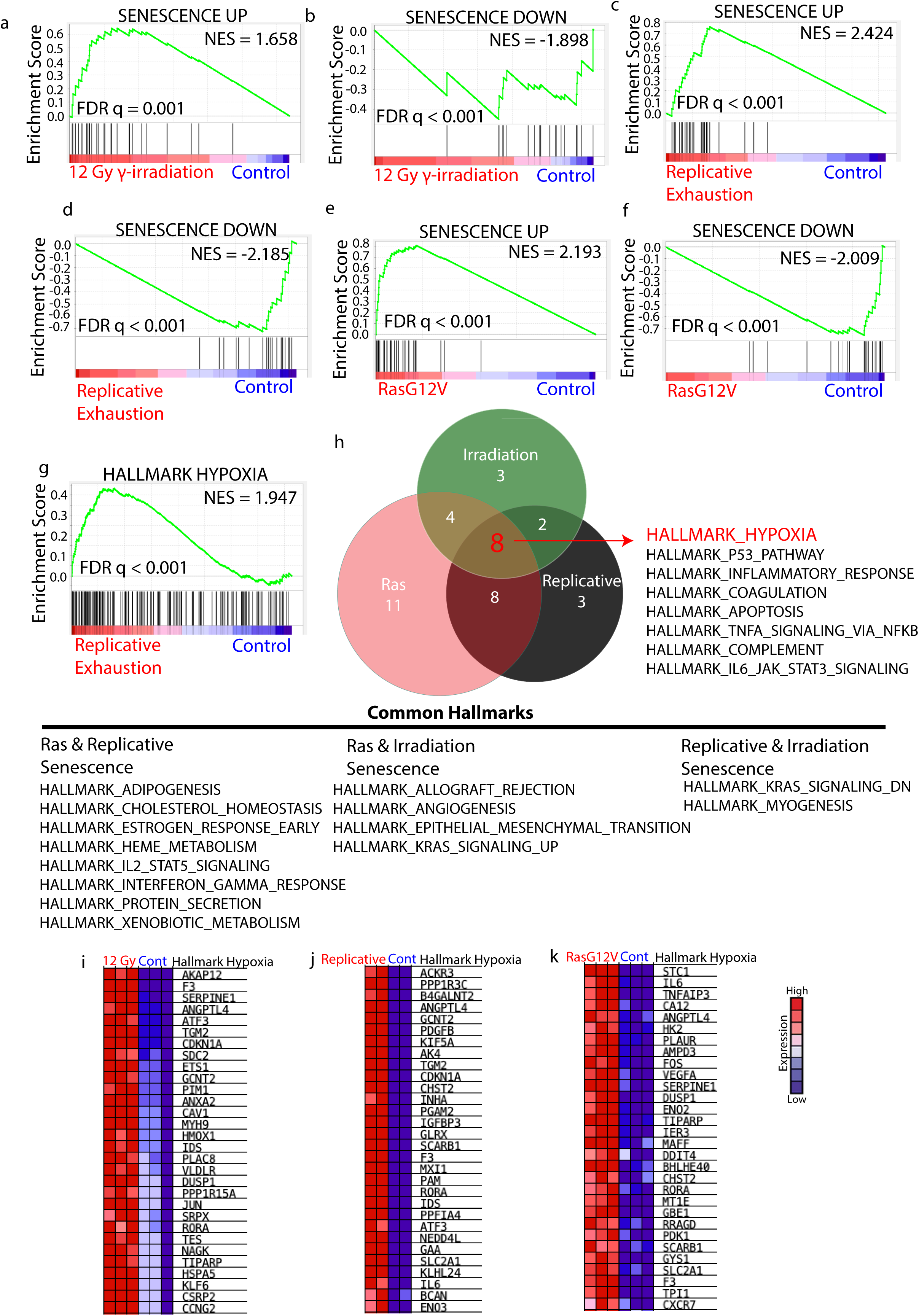
The hallmark hypoxia genes are induced in cellular senescence. (a,b,c,d,e,f) GSEA shows that the senescence up-regulated (a,c,e) and down-regulated (b,d,f) gene sets are induced in the 12 Gy γ-irradiation (a,b), replicative exhaustion (c,d), and RasG12V(e,f) experimental conditions. (g) GSEA shows that the hallmark hypoxia gene set is induced in replicative exhaustion-induced senescence. (h) Eight hallmark expression signatures are commonly upregulated in 12 Gy γ-irradiation, replicative exhausted, and Ras G12V-induced senescent cells. The Venn diagram for the indicated comparisons shows the hallmark expression signatures overlap from all experimental conditions. The bottom panel is a table of the hallmark expression signatures with at least an FDR of q < 0.05 shared between Ras & replicative-exhaustion, Ras & irradiation, and replicative-exhaustion & irradiation-induced senescence, respectively. (i,j,k) Heatmaps showing the leading-edge enrichment of the top hallmark hypoxia 30 genes in (i) 12 Gy γ-irradiation, (j) replicative exhaustion, and (k) RasG12V-induced cells. All GSEA data are from n = 3 independent experiments, except for (all replicative exhaustion related data, j), which is from n = 2 independent experiments.

We then used GSEA to identify other transcriptional signatures besides senescence, which are commonly activated in γ-irradiation, replicative exhaustion, and constitutive-active Ras oncogene-induced senescence. We employ the hallmark gene sets from the Molecular Signatures Database (MSigDB)(Subramanian *et al*., 2005; Liberzon *et al*., 2011; Liberzon *et al*., 2015). Hallmark gene sets summarize and represent specific, well-defined biological states or processes and display coherent expression. These gene sets were generated by a computational methodology based on identifying gene set overlaps and retaining genes that display coordinate expression, thus reducing data noise and redundancy. Strikingly, the hypoxia hallmark was identified as a common expression signature profile from our GSEA of replicative exhausted, γ-irradiated, and Ras-induced cells (Fig. 1g,h). The hallmark hypoxia signature was ranked 3^rd^ and 8^th^ in replicative and Ras-induced senescence, respectively. The hallmark hypoxia signature was ranked 17^th^ in our irradiated cells, ahead of the p53 hallmark response - a key feature of cellular senescence. In the hallmark hypoxia GSEA leading-edge subset from γ-irradiated, replicative exhausted, and constitutive active Ras-induced, the top 30 genes include well-validated HIF-1α targets such as ANGPTL4, PDGFB, and VEGFA (Fig. 1i,j,k). Other hallmark gene sets that were prominent from our GSEA include the inflammatory response, NFκB activation, and IL2/STAT5 signaling (Fig. 1h). These immune-associated expression programs are commonly induced in senescence(Lasry and Ben-Neriah, 2015).

Although HIF-1α pathway activation in SASP is a new finding(Wu *et al*., 2023a), the mechanisms that link senescence triggers, such as Ras and γ-irradiation, to HIF-1α have long been well documented. Indeed, the senescence triggers-Ras and γ-irradiation can activate signalling pathways leading to HIF-1α stabilisation and activation. An avenue for HIF-1α activation by Ras signaling is through NFκB activation(Kang *et al*., 2015). Accordingly, constitutively active mutant Ras can activate NFκB(Chien *et al*., 2006), which induces HIF-1α activation and mRNA transcription(Cummins *et al*., 2006; Rius *et al*., 2008; Fitzpatrick *et al*., 2011). For γ-irradiation, irradiated tumors become reoxygenated, leading to (1) nuclear accumulation of HIF-1α in response to reactive oxygen and (2) enhanced translation of HIF-1α-regulated transcripts secondary to stress granule depolymerization(Moeller *et al*., 2004). Other senescence-inducing stresses, such as oxidative stress(Lopez-Otin *et al*., 2013), are also known to activate HIF-1α(Chandel *et al*., 2000).

How does HIF-1α function in physiology? HIF-1α is a critical sensor of low oxygen levels(Semenza, 1998; Iommarini *et al*., 2017); its regulation in normoxia and hypoxia is as follows: In normoxia, prolyl hydroxylases (PHDs) hydroxylate HIF-1α, triggering von Hippel–Lindau-mediated ubiquitination and proteasomal degradation. Conversely, hypoxia inhibits PHDs and stabilizes HIF-1α, which then translocates into the nucleus and dimerizes with constitutively expressed HIF-1β, activating the HIF complex and triggering the transcription of genes promoting inflammation and cancer(Ruas and Poellinger, 2005).

### HIF-1α target genes are induced in SASP and replicative exhaustion-induced senescence but not in cellular quiescence

We next study the relevance of HIF-1α activation in cellular senescence with a HIF-1α activator Dimethyloxalylglycine, N-(Methoxyoxoacetyl)-glycine methyl ester (DMOG)(Wu *et al*., 2023a). DMOG promotes the steady-state accumulation of HIF-1α and its activation by inhibiting the EGLN/PHD prolyl hydroxylases, which mark HIF-1α for proteasomal degradation(Laitala and Erler, 2018). We performed RNA sequencing of MCF10A cells after overnight treatment with 1 μM of DMOG, with DMSO-treated cells as control. We find that expression of SASP factors, in general, is induced in DMOG-treated cells (Fig. 2a,b). Common SASP factors such as IGFBP3, IL8, and VEGF are among the top 20 genes in the SASP GSEA leading-edge subset (Fig. 2b).

**Figure 2.**
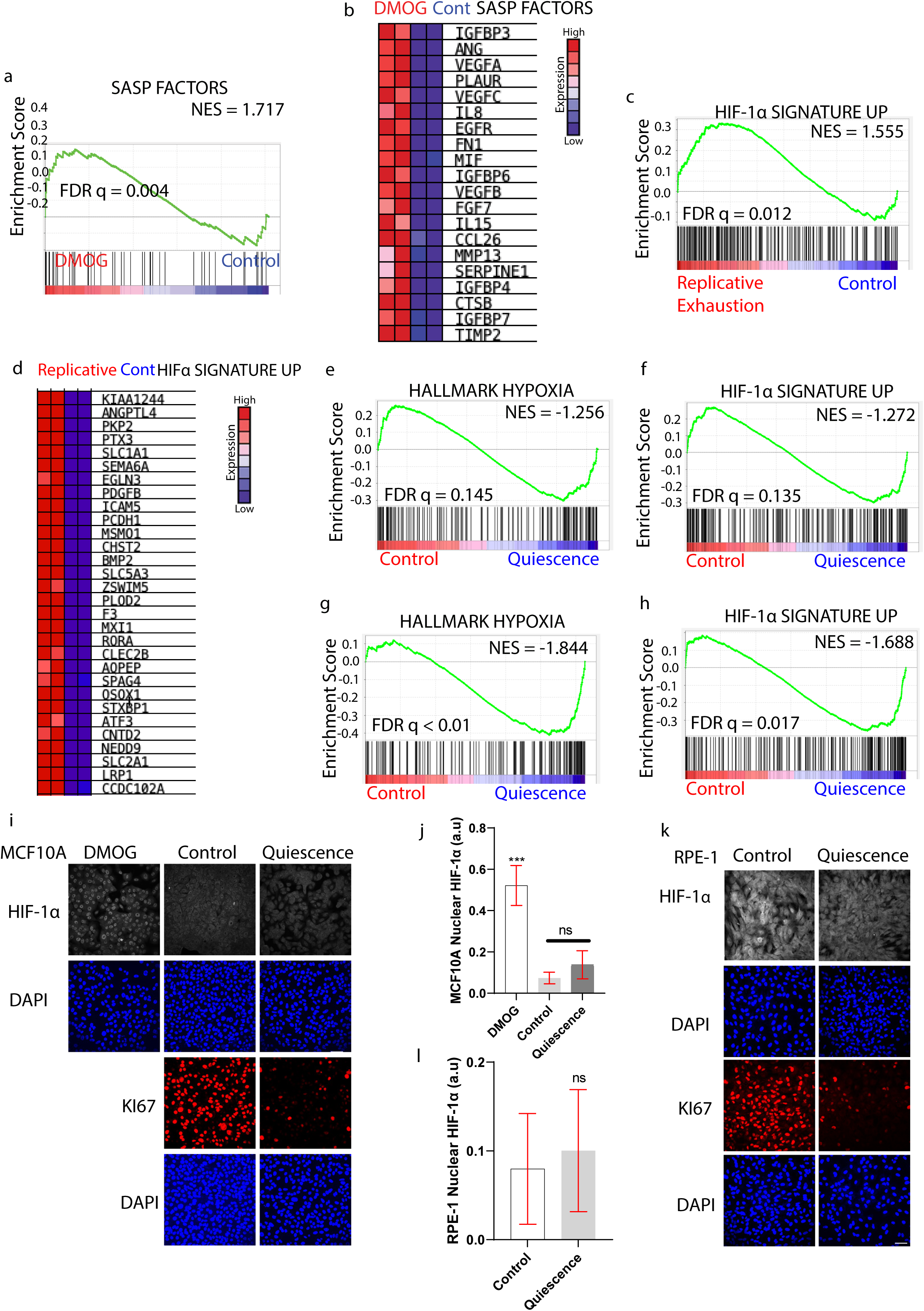
HIF-1α target genes are induced in SASP and replicative exhaustion-induced senescence but not in cellular quiescence. (a) GSEA showing that the SASP factors gene set is enriched in DMOG-treated cells. (b) Heatmap showing the leading-edge enrichment of the top 20 SASP factors in DMOG-treated MCF10A cells. (c) GSEA showing that the custom HIF-1α signature up is enriched in replicative exhausted cells. (d) Heatmap showing the leading-edge enrichment of the top custom HIF-1α signature upregulated 30 genes in replicative exhausted cells. (e,f,g,h) GSEA showing that the (e,g) hallmark hypoxia and (f,h) custom HIF-1α signature up gene sets are not induced in (e,f) IMR90(Tasdemir *et al*., 2016) and significantly suppressed in (g,h) LFS MDAH041 fibroblast cells(Purcell, Kruger, and Tainsky, 2014) undergoing quiescence. (i,j,k,l) HIF-1α is not induced in quiescent MCF10A and RPE-1 cells. Representative images (i,k) and quantitative analysis (j,l) of nuclear HIF-1α and Ki67 in MCF10A(i,j) and RPE-1 cells (k,l). GSEA for (a,c,e,f,g,h) are from n = 3 independent experiments. Data for (b) and (d) are from n = 2 independent experiments. Data for (j) and (l) are means ± SD from n = 6 fields of technical replicates (∼218 cells per field) and n = 7 fields (∼254 cells per field) from 2 independent experiments respectively. ns, not significant; analysed with Mann-Whitney test. Scale bars, 5 μm.

We confirm HIF-1α activation in our transcriptomes of senescent cells with a HIF-1α signature gene set. The HIF-1α signature gene set was constructed by including genes that were upregulated by DMOG in DESeq analysis (*p*(adjusted) < 0.05, fold change ≥4). DESeq analyzes RNA-seq data to identify differentially expressed genes, accounting for overdispersion and library size normalization. The custom HIF-1α activation gene set is significantly enriched in replicative exhaustion-induced senescent cells (Fig. 2c,d). Notably, we previously showed that HIF-1α activation expression signature is activated in γ-irradiation, Ras-induced, and centrosome amplification-induced senescent cells transcriptomes(Wu *et al*., 2023a). Senescence is a permanent, stress-induced cell cycle arrest associated with a secretory phenotype. In contrast, quiescence is a reversible, adaptive response to cellular stress in which cells enter a resting state. We asked whether HIF-1α is specifically activated in cellular senescence but not in quiescence. To that end, we analyze two previously published datasets for the induction of cellular quiescence in IMR90 fibroblasts(Tasdemir *et al*., 2016) (Fig. 2e,f) and LFS MDAH041 fibroblast cells(Purcell, Kruger, and Tainsky, 2014) (Fig. 2g,h). Our analysis revealed that both the hallmark hypoxia and HIF-1α activation gene sets were either not induced (Fig. 2e,f) or significantly suppressed (Fig. 2g,h). Considering that different cell lines have different genetic backgrounds, we further analyzed HIF-1α activation during quiescence on our experimental cell lines RPE-1 and MCF10A cells as well. Accordingly, we observed that nuclear HIF-1α is not induced by cellular quiescence in both MCF10A (Fig. 2i,j) and RPE-1cells (Fig. 2k,l). Thus, our data demonstrates that HIF-1α is not activated in quiescence.

### Centrosome amplification induces hallmarks of cellular senescence in MCF10A epithelial cells and WI38 fibroblasts

We next shift our focus from the conventional stresses that induce senescence to analyse the consequences of centrosome amplification-induced senescence(Wu *et al*., 2023a). Centrosome amplification was induced in cells by overexpressing Polo-like kinase 4 (PLK4), the master regulatory kinase for centrosome duplication(Bettencourt-Dias *et al*., 2005; Habedanck *et al*., 2005). PLK4 was transiently induced, and subsequent analysis was performed at time points where increased PLK4 mRNA expression was no longer detectable(Godinho *et al*., 2014). In most experiments described below, cells with the transient overexpression of a truncated PLK4 mutant (Control PLK4^1-608^) with kinase activity, which did not localize to the centrosomes(Guderian *et al*., 2010), were used as the negative control.

Retinal Pigment Epithelial-1 (RPE-1) and MCF-10A epithelial cells were previously reported to enter senescence seven days post-induction with extra centrosomes(Wu *et al*., 2023a). Whether cell types besides epithelial cells enter senescence when induced with centrosome amplification remains unclear. WI38 normal lung fibroblasts, which have the normal number of centrosomes (Fig. 3a,b), is a common cell line model for studying senescence. Here, we find that WI38 lung fibroblasts also exhibit senescence-associated beta-galactosidase activity (∼60% positive cells, Fig. 3c,d) seven days post extra centrosomes induction by transient PLK4 expression (Fig. 3a,b).

**Figure 3.**
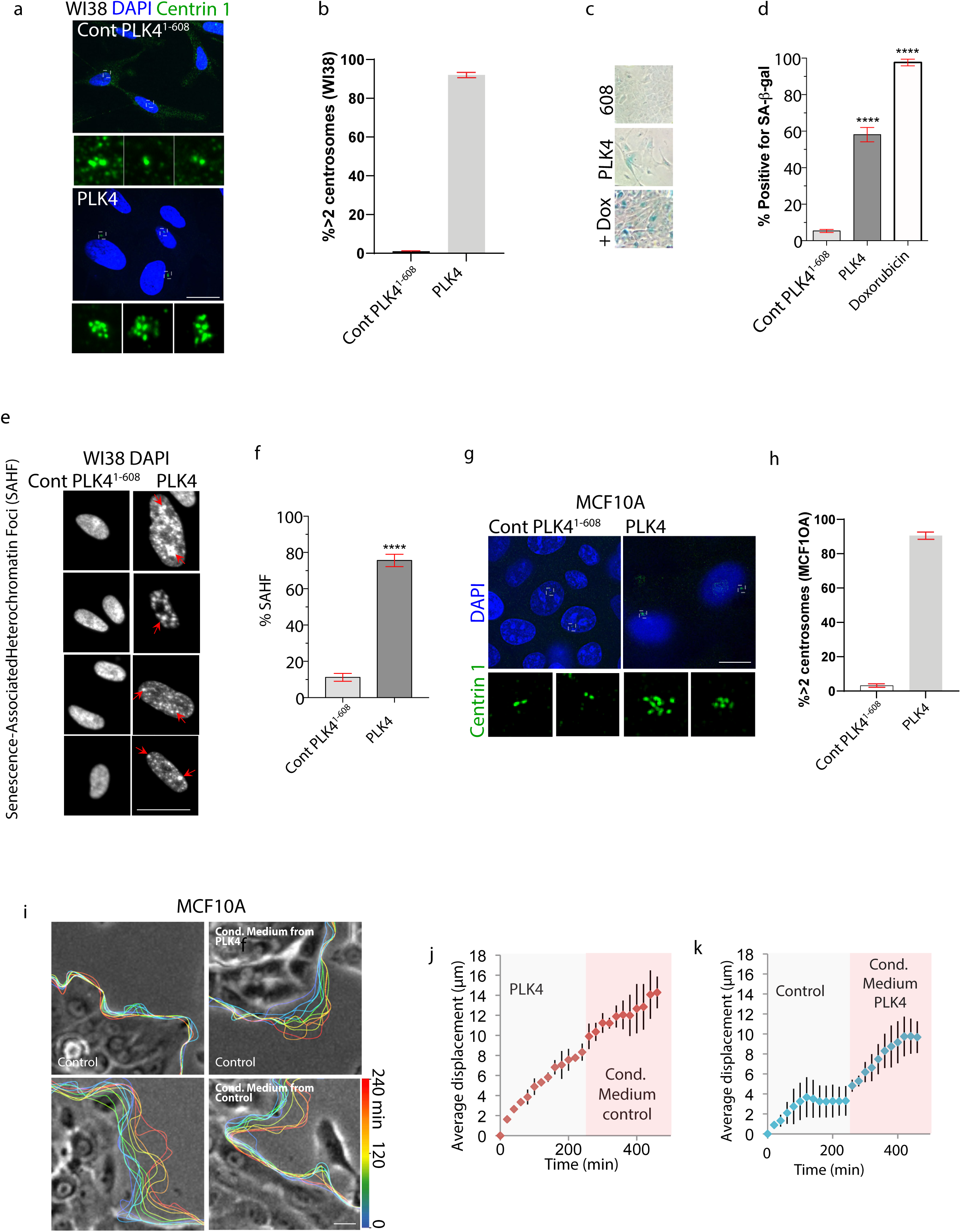
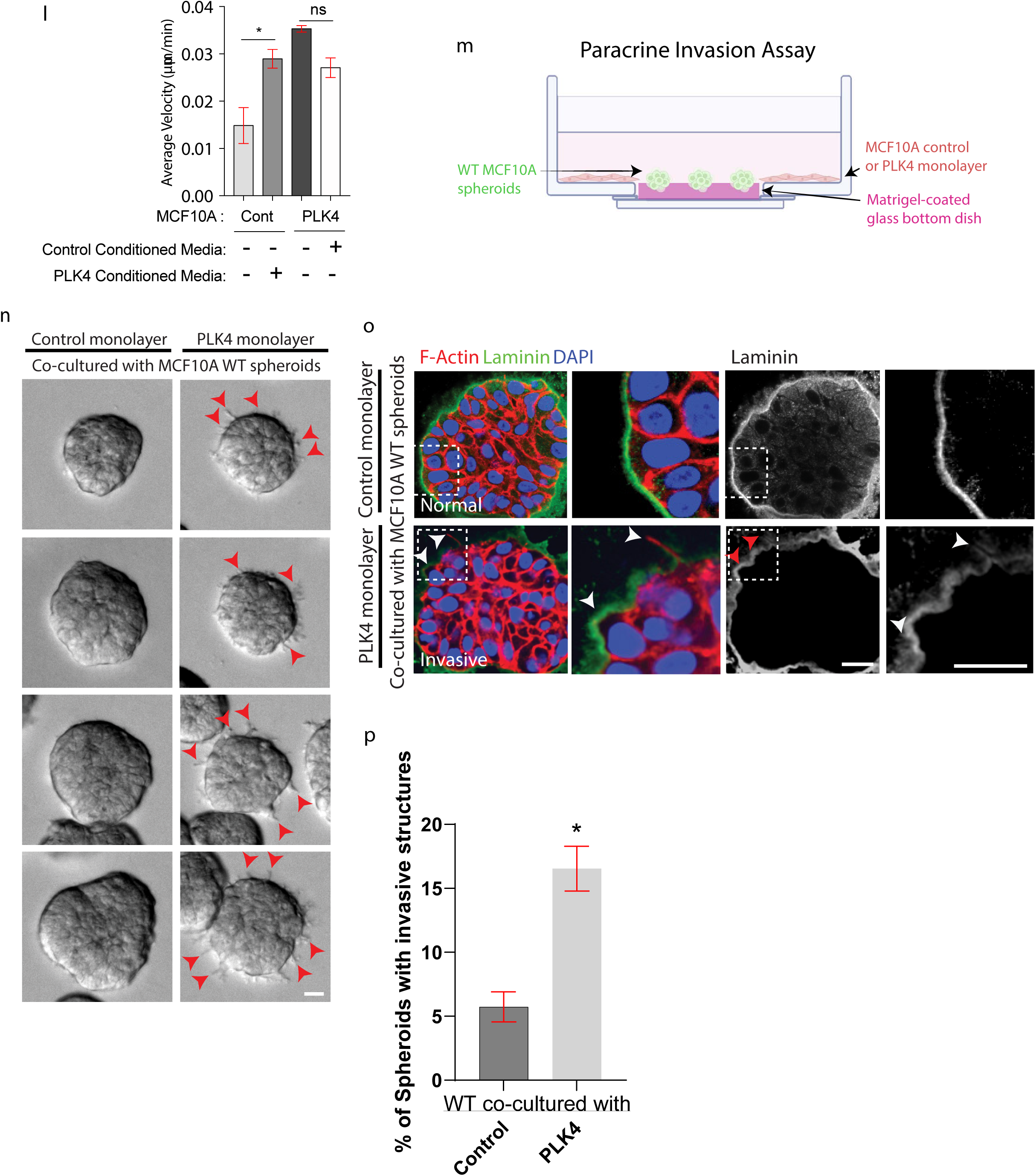
Centrosome amplification induces hallmarks of cellular senescence in MCF10A epithelial cells and WI38 fibroblasts. (a, b) Fraction of Control PLK4^1-608^ and PLK4-induced WI-38 cells with more than two centrosomes was determined by immunostaining of centrin-1. Representative images (a) and quantification (b) of WI38 cells with (PLK4) and without (Control PLK4^1-608^) centrosome amplification. Scale bars, 5 μm. Data are means ± SEM from n=3 independent experiments. (c,d) Increased SA-β-Gal staining in WI38 fibroblasts with centrosome amplification. Representative images (c) and quantification (d) of SA-β-Gal (blue) staining of WI38 fibroblasts with and without centrosome amplification or positive control with doxorubicin treatment. (e,f) Centrosome amplification induces Senescence-Associated Heterochromatin Foci formation in WI38 cells visible with Hoechst. Representative images (e) and quantification (f) of senescence-associated heterochromatin foci (Hoechst) in WI38 cells with and without centrosome amplification. (g,h) Fraction of Control PLK4^1-608^ and PLK4-induced MCF10A cells with more than two centrosomes was determined by immunostaining of centrin-1. Representative images (g) and quantification (h) of MCF10A cells with (PLK4) and without (Control PLK4^1-608^) centrosome amplification. Scale bars, 5 μm. Data are means ± SEM from n=3 independent experiments. (i-l) Conditioned media from MCF10A cells with centrosome amplification stimulates the leading-edge displacement of wild-type cells. (i) Representative images of leading-edge displacement in cells with and without centrosome amplification, with or without conditioned media swap. The colour code on the heatmap and images represents the position of the leading edge at different time points. (j,k) Quantitative analysis of leading-edge displacement in MCF10A cells with (PLK4, j) or without (control, k) centrosome amplification, before and after exchange with the indicated conditioned medium. (l) Conditioned media from MCF10A cells with centrosome amplification significantly increases the average protrusion velocity of control cells. Shown is the average velocity of leading-edge displacement in cells with and without centrosome amplification, with or without conditioned media swap. (m,n) Paracrine factors induced co-culture invasion assay with and without centrosome amplification-induced senescence cells. Schematic diagram (m) and representative images (n) of WT MCF10A spheroids co-cultured with control (left) or centrosome-amplified PLK4 senescence cell monolayer (right). Red arrows indicate invasive protrusions of WT MCF10A spheroids. (o) Representative images of MCF10A WT spheroids co-cultured with MCF10A monolayer cells with (PLK4) and without (Control) centrosome amplification immunostained for F-Actin (red), Laminin (green), and DAPI (blue). (p) Quantification of WT MCF10A spheroids with invasive structures when co-cultured with (PLK4) and without (Control) centrosome amplification. Data are means ± s.e.m. from n = 4 independent experiments. Data for (b,d,f,h,l) are means ± SEM from n = 3 independent experiments from at least 79 cells for (b), 90 cells for (d), 75 cells for (f), 85 cells for (h), and 42 cells for (l) for each experimental group. Data for (p) is means ± SD from n = 4 groups of technical replicates (∼29 spheroids per group) from 2 independent experiments. *P < 0.05,****P <0.0001; ns, not significant; analysed with Student’s t-test (b,d,f,h,l) and Mann-Whitney test (p). Scale bars, 5 μm.

Additionally, senescence-associated heterochromatin foci (SAHF) were observed in Hoechst-stained WI38 fibroblasts after the induction of centrosome amplification by transient PLK4 expression (Fig. 3e,f). SAHF observed with Hoechst staining has been previously reported as a cell-type-specific phenotype(Kosar *et al*., 2011). Indeed, such a striking Hoechst labelled SAHF seen in WI38 cells is not obvious in MCF10A cells. Even with HP1γ staining for SAHF, used previously(Nader *et al*., 2021), the localization of senescence-associated heterochromatin foci in MCF10A cells is harder to discern (Supplementary Fig. 1a).

The induction of cellular motility by secreted factors is a hallmark of SASP(Coppe *et al*., 2010). MCF10A epithelial cells are an excellent model for studying cell motility in 2D and 3D cultures. Since the majority of MCF10A cells have normal numbers of centrosomes (Fig. 3g,h), they are also ideal for studying the effects of centrosome amplification. We previously found that centrosome amplification can induce cellular invasion induced by paracrine factors during SASP(Wu *et al*., 2023a). Cell protrusions are dynamic, outward extensions of the plasma membrane that play a crucial role in processes like cell migration, adhesion, and sensing the environment. To visualize whether cells with centrosome amplification can induce protrusions of normal cells in a paracrine manner, we swapped the conditioned medium between MCF10A cells with centrosome amplification and the conditioned medium from control cells without centrosome amplification (Fig. 3g,h). As expected, the control conditioned medium did not alter the protrusion velocity of cells’ migratory front with centrosome amplification (Fig. 3i,j,l, and Supplementary Video 1). By contrast, conditioned media from cells with centrosome amplification induced an approximately 2-fold increase in the protrusion velocity of the migratory front in cells lacking centrosome amplification (Fig. 3i,k,l and Supplementary Video 2). To better assess invasion, we designed an MCF10A acini and centrosome amplification cells co-culture assay (Fig. 3m). Our co-culture assay revealed that centrosome amplification-induced senescence cells promote the invasiveness of WT MCF10A acini in a paracrine manner (Fig. 3n-p, Supplementary Video 3). Notably, invasive protrusions of WT MCF10A spheroid penetrated the laminin-rich basement membrane (Fig. 3o), induced by cells with centrosome-amplification in a paracrine manner, which is a hallmark of SASP.

### Centrosome amplification induced a non-canonical SASP with an inflammatory response dominated by HIF-1α rather than the classical NFκB

Our data indicated that centrosome amplification induced a non-canonical SASP(Wu *et al*., 2023a). Although centrosome amplification generated many cellular features and gene expression characteristics of senescence, several aspects of the response were distinct from senescence induced by canonical triggers(Wu *et al*., 2023a). First, the SASP induced by centrosome amplification was distinct from the SASP induced by conventional triggers dominated by NF-κB activation. NF-κB is generally thought to be the primary determinant of the SASP secretome(Kang *et al*., 2015), and DNA damage signaling was recently shown to activate NF-κB through the transcription factor GATA4(Rodier *et al*., 2009; Kang *et al*., 2015). However, there is no detectable GATA4 mRNA in MCF10A cells, with or without centrosome amplification (Fig. 4a). Additionally, centrosome amplification induction of proinflammatory cytokines was weaker by comparison with the SASP induced by γ-irradiation. This is because we can observe an enrichment of various inflammatory expression signatures, including interferon alpha, gamma, and complement activation, in MCF10A cells with γ-irradiation-induced senescence compared to centrosome amplification-induced senescence cells (Fig. 4b,c,d)(Acosta *et al*., 2013; Kang *et al*., 2015), thus suggesting a relatively weaker inflammatory response induced by centrosome amplification.

**Figure 4.**
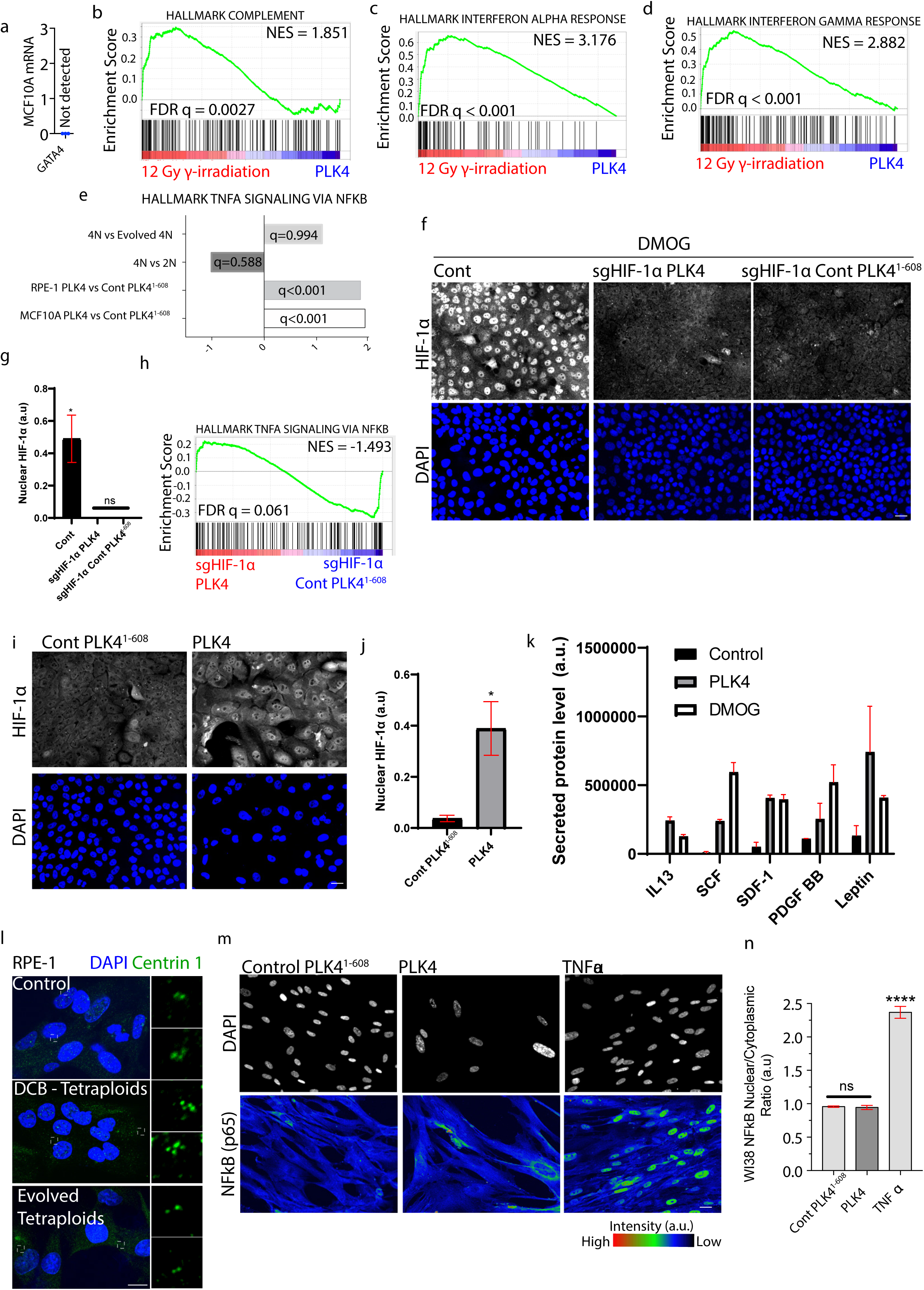
Centrosome amplification induces an inflammatory response dominated by HIF-1α rather than a SASP controlled by NFκB. (a) Undetectable levels of GATA4 in MCF10A cells before and after induction of centrosome amplification. Data are from n = 3 independent experiments (b,c,d) GSEA shows that inflammatory genes from complement, interferon alpha and interferon-gamma response are more significantly enriched in 12 Gy γ-irradiated induced senescent cells than senescent cells induced by centrosome amplification. (e) Induction of NF-κB regulated genes after centrosome amplification from PLK4 transient overexpression, but not from cytokinesis failure. Shown are GSEA FDR: q values and normalized enrichment score (NES) for the indicated cell lines. (f,g) CRISPR-mediated gene disruption of HIF1A abolishes the induction of HIF-1α nuclear staining by DMOG. Representative immunofluorescence images (f) and quantification (g) of HIF-1α and DAPI in DMOG treated Control, sgHIF-1α PLK4 and sgHIF-1α Control PLK4^1-608^. (h) CRISPR-mediated gene disruption of HIF1A abolishes the induction of NF-κB regulated genes by centrosome amplification. GSEA revealed a statistically insignificant suppression of NF-κB-regulated genes in sgHIF1A cells with centrosome amplification relative to sgHIF1A control cells. (i,j) Induction of nuclear HIF-1α after centrosome amplification-induced senescence. Representative images (i) and quantification (j) of nuclear HIF-1α in MCF10A cells with or without centrosome amplification. (k) Quantification of HIF-1α activation targets. Protein levels of secreted factors IL13, SCF, SDF-1, PDGF BB, and Leptin in Control, centrosome amplification-induced senescence PLK4, and DMOG treated Control cells. (l) RPE-1 tetraploids and evolved tetraploids with extra centrosomes and the normal number of centrosomes, respectively, were determined by immunostaining of centrin-1. Representative images of RPE-1 diploids, tetraploids and evolved tetraploids. (m,n) Lack of NF-κB nuclear accumulation after centrosome amplification. (m) Representative images of p65 RELA (NF-κB) and DAPI (grey) in WI38 cells with and without centrosome amplification as compared with TNFα treated cells. (n) Quantification of p65 RELA (NF-κB) nuclear/cytoplasmic ratio in WI-38 cells with or without centrosome amplification compared to TNFα treatment. GSEA for (b,c,d,e,h) are from n = 3 independent experiments. Data for (g,j,n) are means ± SEM from n = 4 independent experiments from at least 147 cells for (g), at least 58 cells for (j), and at least 28 cells for (n) per experimental group. ns, not significant, ****P <0.0001; analysed with Mann-Whitney test. Scale bars, 5 μm.

By GSEA, we observed enrichment for NF-κB-regulated genes in the samples where centrosome amplification was induced by PLK4 transient overexpression (Fig. 4e). However, this enrichment for NF-κB-regulated genes was abolished by CRISPR-mediated gene disruption of HIF-1α (Fig. 4e,f,g), suggesting that the shared gene targets of HIF-1α with NF-κB contribute to the enrichment of NF-κB-regulated genes(Palazon *et al*., 2014). We also confirmed that HIF-1α is activated in these centrosome amplification-induced MCF10A senescent cells, as we observed an increase in both the nuclear localization of HIF-1α (Fig. 4h) and increased secreted factors which are targets of HIF-1α activation (Fig. 4i)(Grosfeld *et al*., 2002; Ceradini *et al*., 2004; Schito *et al*., 2012; Gao *et al*., 2015; Chiba, Ohnishi, and Matsuguchi, 2025).

As an orthogonal approach to examine the effects of centrosome amplification, we employed a PLK4 expression-independent approach to generate extra centrosomes. We compared the transcriptomes of newly generated tetraploid cells from dicytochalasin B-induced cytokinesis failure with doubled centrosome content to those of the control cells (Fig. 4j). Both parental diploid cells and “evolved” tetraploid cells that had spontaneously lost their extra centrosomes were used as controls (Fig. 4j)(Ganem, Godinho, and Pellman, 2009; Ganem *et al*., 2014). Specifically, we did not observe a significant enrichment for NF-κB-regulated genes in RPE-1 cells where centrosome amplification was induced by cytokinesis failure (Fig. 4e). Strikingly, quantitative imaging indicated that centrosome amplification in WI38 lung fibroblasts did not result in any detectable increase in the nuclear localization of NF-κB (Fig. 4k,l). In contrast, our positive control TNFα (activator of NF-κB) treated cells showed a robust NF-κB nuclear localization. Thus, centrosome amplification induces a non-canonical SASP dominated by HIF-1α activation.

### Centrosome amplification-generated micronuclei do not induce the cGAS/Sting mediated interferon response in SASP

Second, the effects of centrosome amplification are distinct from those of aneuploidy. Although aneuploidy can induce cell cycle arrest and senescence, it is established that this occurs at high levels of aneuploidy that disrupt DNA replication and cause DNA damage(Santaguida *et al*., 2017; Soto *et al*., 2017). Accordingly, we confirm that low-dose MPS1 inhibition, which is known to induce moderate levels of chromosome missegregation and micronuclei induction at comparable levels of aneuploidy(Godinho *et al*., 2014) and micronuclei generated to that induced by centrosome amplification, does not trigger proliferation arrest (Fig. 5a,b,c) or trigger HIF-1α nuclear translocation (Fig. 5d,e). However, for cells to be defined as senescent, they should at least display proliferation arrest together with other canonical features of senescence(Perez-Mancera, Young, and Narita, 2014; Gorgoulis *et al*., 2019). Our data indicate that moderate levels of chromosome missegregation and micronuclei induction do not induce cellular senescence.

**Figure 5.**
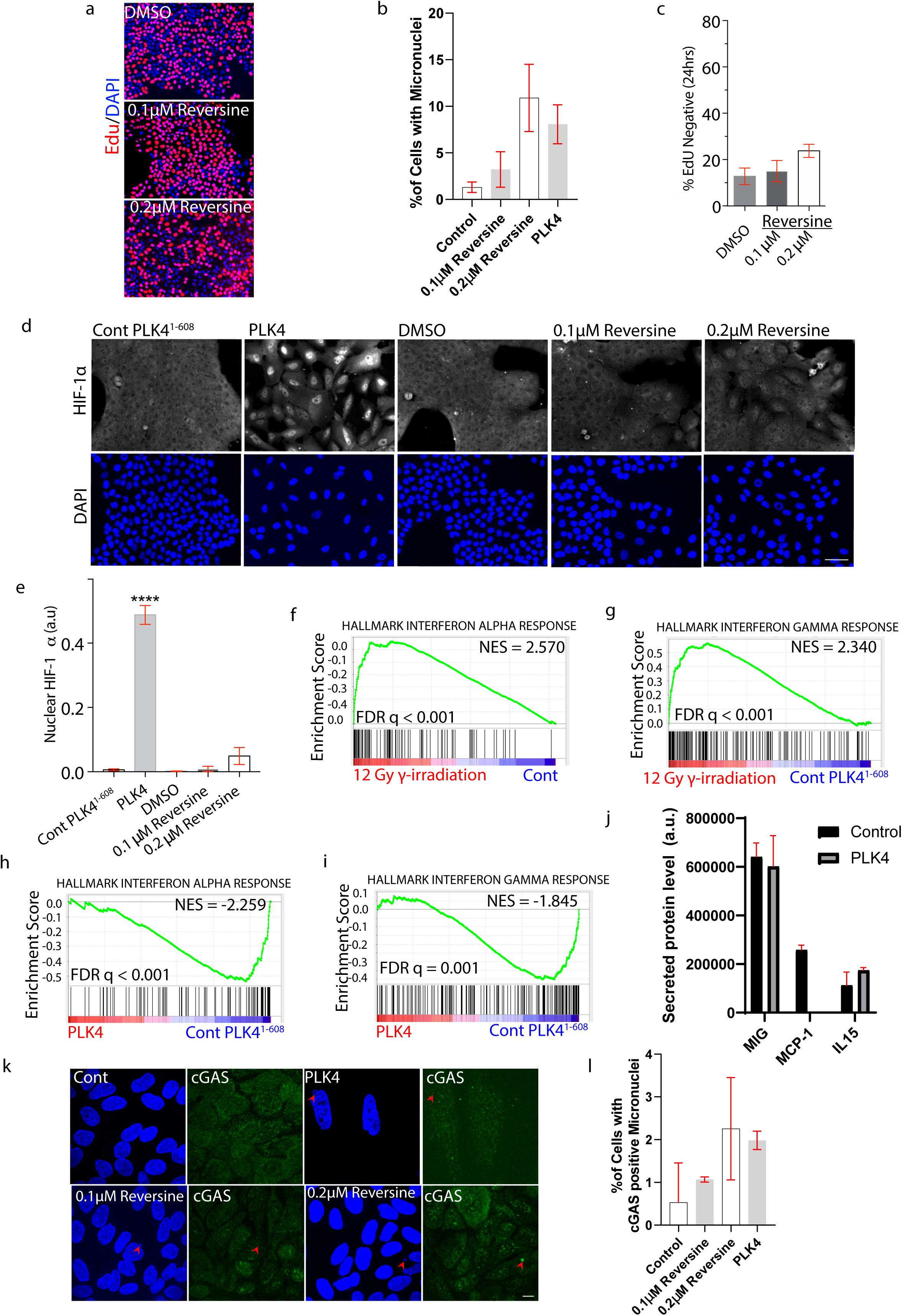
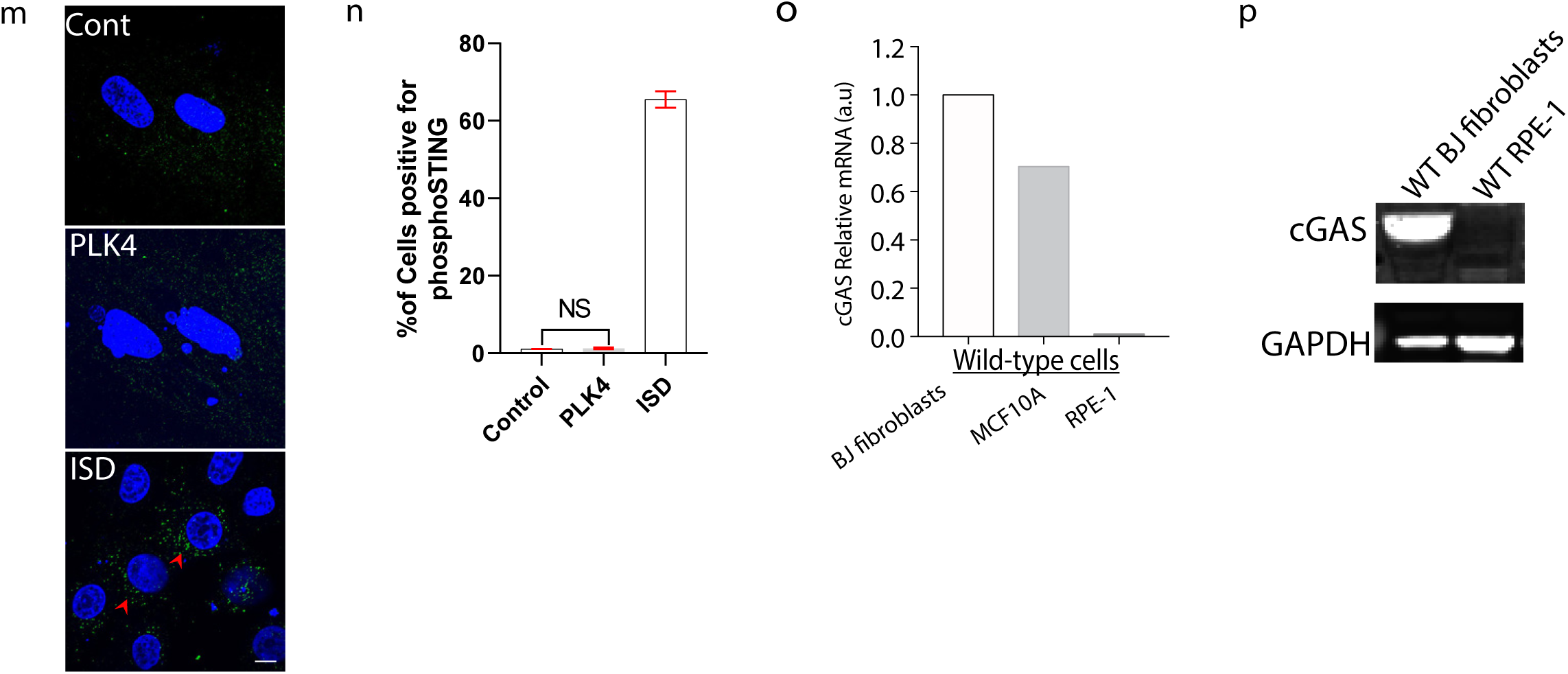
Centrosome amplification generated micronuclei do not induce the cGAS/Sting mediated interferon response in SASP. (a,b,c) Low-dose MPS inhibition by reversine at concentrations that generate comparable levels of micronuclei to centrosome amplification from transient PLK4(Godinho *et al*., 2014)does not induce proliferation arrest. Representative images (a) and quantification (b) of micronuclei (c) EdU (red) and DAPI (blue) in MCF10A cells with (PLK4) and without (Control PLK4^1-608^) centrosome amplification or control cells treated with 0.1 μM or 0.2μM reversine for 24 hours. (d,e) Low-dose MPS inhibition by reversine, at concentrations that generate comparable or higher levels of aneuploidy than centrosome amplification from transient PLK4(Godinho *et al*., 2014), does not induce nuclear HIF-1α. Representative images (d) and quantification (e) of nuclear HIF-1α levels in MCF10A cells with (PLK4) and without (Control PLK4^1-608^) centrosome amplification or wild-type cells treated with either 0.1μM or 0.2μM reversine. (f,g) Induction of interferon-alpha (f) and interferon-gamma (g) response genes in cells after γ-irradiation by GSEA. (h,i) Suppression of interferon-alpha and interferon-gamma response expression in cells with centrosome amplification. GSEA revealed the downregulation of genes annotated as Hallmark interferon alpha response (h) and Hallmark interferon-gamma response (i) in MCF10A cells with centrosome amplification relative to control. (j) Quantification of secreted protein levels of pro-inflammatory cytokines MIG, MCP-1, and IL15 in Control, centrosome amplification-induced senescence PLK4, and DMOG treated Control cells. (k,l) Low levels of cGAS-positive centrosome amplification generated micronuclei. Representative images (k) and quantification of (l) micronuclei positive for cGAS immunostaining. (m,n) Lack of STING Ser366 phosphorylation in perinuclear cytoplasmic regions in cells with micronuclei generated by centrosome amplification. Representative images (m) and quantification of (n) cells with centrosome amplification generated micronuclei positive for perinuclear/cytoplasmic STING Ser366 phosphorylation immunostaining. (o) Undetectable levels of cGAS in RPE-1 cells. Relative mRNA level of cGAS in BJ, MCF10A, and RPE-1 cells by RT-qPCR. (p) cGAS protein is undetectable in RPE-1 cell lysates. cGAS and GAPDH (loading control) immunoblots of lysates from BJ and RPE-1 cells. Data for (b,l) are means ± SEM from n = 3 independent experiments from at least ∼157 cells for (b) and at least ∼70 micronuclei for (l) per experimental group. Data for (c,e,n) are means ± SEM from n = 8 fields for (c) with ∼142 cells per field, n = 4 fields for (e) with ∼121 cells per field, and n = 19 fields for (n) with ∼8 cells per field from 2 independent experiments, respectively. GSEA for (f,g,h,i) are from n = 3 independent experiments. ****P <0.0001; analysed with Student’s t-test (b,l) and Mann-Whitney test (c,e,n). Scale bars, 5 μm.

Lastly, senescence from various conventional triggers, for example, 12 Gy γ-irradiation of MCF10A (Fig. 5 f,g), triggers an interferon response, possibly due to cytosolic mitochondrial DNA release activating STING(Sato and Hayashi, 2024). However, in the SASP that occurs after centrosome amplification in MCF10A cells, interferon response genes are suppressed, not induced (Fig. 5h,i,j)(Liebler *et al*., 1994; Liu *et al*., 2001; Lee *et al*., 2014). A major consequence of centrosome amplification is the generation of micronuclei by chromosome missegregation. Our data is consistent with reports indicating that micronuclei do not activate cGAS-mediated interferon response(Flynn, Koch, and Mitchison, 2021; Sato and Hayashi, 2024; Takaki *et al*., 2024). Indeed, we observed a low level of cGAS and Sting activation, with 2% of micronuclei that are cGAS positive as compared to control (Fig. 5k,l). Immunostaining of MCF10A cells transfected with ISD (IFN stimulatory DNA) dsDNA revealed robust perinuclear cytoplasmic phosphorylation of STING on Ser366. By contrast, centrosome amplification cells containing micronuclei failed to induce cytoplasmic STING phosphorylation (Fig. 5m,n).

Moreover, RPE-1 cells, which manifest an upregulation of secreted protein expression after senescence induction by various triggers(Santaguida *et al*., 2017), including centrosome amplification(Wu *et al*., 2023a), lack detectable expression of cGAS with immunoblotting (Fig. 5o,p). The cGAS expression-positive control BJ fibroblasts were known to have an intact cGAS–STING pathway(Flynn, Koch, and Mitchison, 2021). The lack of cGAS expression in RPE-1 cells was also reported previously by other laboratories(Basit *et al*., 2020; Nader *et al*., 2021). Thus, consistent with a previous report(Nader *et al*., 2021), our data suggest that cGAS is not essential for cellular senescence in RPE-1 cells.

In summary, we have revealed that HIF-1α target genes are specifically induced in senescence but not in quiescence. Consistent with our previous finding(Wu *et al*., 2023a), we have further demonstrated how centrosome amplification induces a non-canonical SASP dominated by HIF-1α activation, rather than the classical NFκB signaling. Lastly, we reveal that during SASP, centrosome amplification-generated micronuclei do not activate the cGAS/Sting-mediated interferon response. Thus, we conclude that SASP constitutes HIF-1α activation but is independent of the micronuclei-induced cGas activation.

---

**Supplementary Figure 1.**
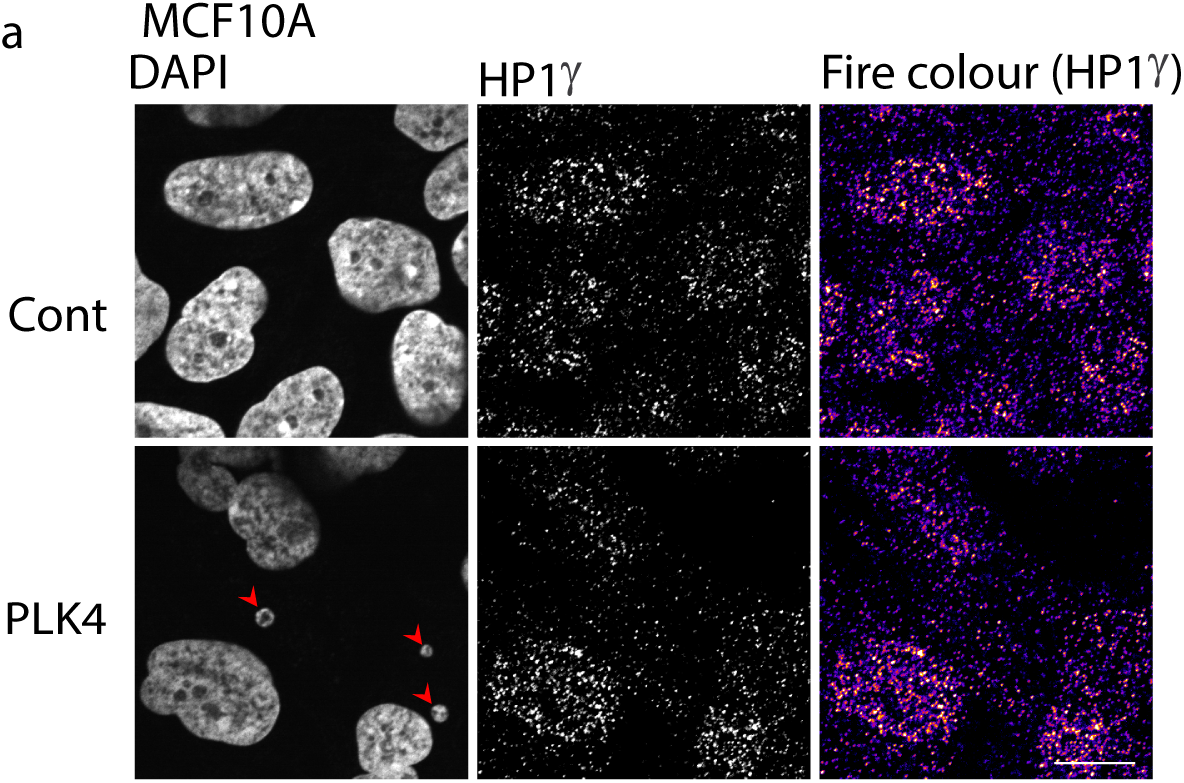
(a) Senescence-associated heterochromatin foci in MCF10A cells is not obvious with HP1γ immunostaining. Representative images from 4 independent experiments, of MCF10A cells with (PLK4) and without (Control) centrosome amplification. Red arrows indicate micronuclei. Scale bars, 5 μm.

**Supplementary Video 1. Conditioned media from control cells does not affect the leading edge movement of cells with centrosome amplification.** Live imaging of MCF10A cells with centrosome amplification before (left) or after (right) control conditioned media swap.

**Supplementary Video 2. Conditioned media from MCF10A cells with centrosome amplification stimulates the leading edge movement of control cells.** Live imaging of control MCF10A cells without centrosome amplification before (left) or after (right) centrosome amplification conditioned media swap.

**Supplementary Video 3. Co-culture assay with centrosome amplification-induced senescence cells promote invasiveness of WT MCF10A spheroids in a paracrine manner.** Live imaging of WT MCF10A spheroids co-cultured with control monolayer without centrosome amplification (left) or centrosome-amplified PLK4 monolayer (middle, right). Red arrows indicate invasive protrusions of WT MCF10A spheroids.

## Conflict of Interest

The authors declare that the research was conducted without any commercial or financial relationships that could be construed as a potential conflict of interest.

## Acknowledgments

We dedicate this work posthumously to J.Campisis for her leadership in SASP discovery and to Angelika Amon for her interest in our work. We gratefully thank D.Pellman, I. Ghobrial, S. Papathanasiou, A. Spektor, A.J. Holland, W. Johnson, Y. Kaplan, A. Yap, S. Liu, K. Briggs, and W. Kaelin for their unstinting support and advice during this project. A.Spektor for γ-irradiation experiments, L. Hodgson and H. Zhang for technical advice and reagents, Y. Wang, D. Deconti, and B. Lawney from the Centre for Cancer Computational Biology and X. Qiu and H. Long from the Center for Functional Cancer Epigenetics for help with bioinformatic analysis. SKW was supported by a Leukemia and Lymphoma Society Fellow Award (5454-17) and a Young Individual Research Grant (MOH-OFYIRG20nov-0019) by the National Medical Research Council of Singapore. This work was also supported by a Research Scholarship Block Research Fellow Scheme and MOE Tier 1 by the Singapore Ministry of Education to S.K.W. and B.C.L., and National Research Foundation Grant (NRF-MSG02023-0001) to B.C.L. LD was supported by the National Key Research and Development Program of China (2022YFA1302800) and the National Natural Science Foundation of China (32270779). RP was supported by a Susan G. Komen Postdoctoral Fellowship (PDF15333560). We acknowledge the Nikon Imaging Center at Harvard Medical School for imaging support.

## Notes

### Competing Interest Statement

The authors have declared no competing interest.

### Summary of Updates

More experiments have been added with a revised text to become a paper of high rigour.

